# Centrosomal P4.1-associated protein (CPAP) positively regulates endocytic vesicular transport and lysosome targeting of EGFR

**DOI:** 10.1101/2020.08.03.234690

**Authors:** Radhika Gudi, Viswanathan Palanisamy, Chenthamarakshan Vasu

## Abstract

Centrosomal P4.1-associated protein (CPAP) plays a critical role in restricting the centriole length in human cells. Here, we report a novel, positive regulatory role for CPAP in endocytic vesicular transport (EVT) and lysosome targeting of internalized-cell surface receptor EGFR. We observed that higher CPAP levels cause an increase in the abundance of multi-vesicular body (MVB) and EGFR is detectable in CPAP-overexpression induced puncta. While the surface levels, and total and phosphorylated cellular levels of EGFR are higher under CPAP deficiency, ligand-engagement induced internalization of this receptor is not impacted by CPAP levels. Furthermore, routing of EGFR into early endosomes is not influenced by CPAP. However, most importantly, we found that CPAP is required for targeting ligand-activated, internalized EGFR to lysosome. Transport of ligand-bound EGFR from early endosome to late endosome/MVB and lysosome is severely diminished in CPAP-depleted cells. Moreover, CPAP depleted cells showed diminished ability to form MVB structures upon EGFR activation. These observations show a positive regulatory role for CPAP in early endosome to late endosome transport and lysosome targeting of ligand-bound EGFR-like cell surface receptors. Overall, identification of this regulatory role for CPAP in endocytic trafficking of EGFR provides new insights in understanding the cellular functions of CPAP.

## Introduction

Cells can internalize extracellular material, ligands, and surface molecules including receptors by endocytosis. Endocytosis is a fundamental biological process that maintains the cellular, tissue and organismal homeostasis. Internalized molecules traffic through, and are sorted by, a series of tubulovesicular compartments including endocytic vesicles^1^. Vesicular transport pathways play an essential role in the delivery of intra- and extra-cellular cargo, and are critical for cell-to-cell communication^2–6^. Dysregulated cargo delivery pathways can have catastrophic effects and are associated with cancer, neurological diseases, and immunodeficiency^2,7–12^.

Post endocytosis, the cell surface receptors such as epidermal growth factor receptor (EGFR) are routed through different functional stages of endosomes, and can have diverse fates such as: 1) be recycled back to the surface, 2) get targeted for degradation, and/or 3) be released outside the cell in exosomes^13^. Early endosomes (EE) serve as a sorting nexus for, and play a major role in deciding the cellular fate of, endocytosed cargo^1,14^. While the cargo can be recycled back to the surface through recycling endosomes (RE) ^15^, regulatory mechanisms exist to ensure that the cargo destined for degradation is routed to lysosomes^16^. Lysosome targeting has two additional steps: (i) the formation of an intermediate endocytic organelle referred to as the multi-vesicular body (MVB), also called as multi-vesicular endosome (MVE) or the late endosome (LE) ^17,18^, and (ii) the sequential acquisition and activation of GTPases Rab5 and Rab7^7,19,20^. Rab5 to Rab7 conversion is an essential step that marks the maturation of EE to MVBs^21,22^. Each MVB possesses a limiting membrane that encloses several smaller vesicles (referred to as intraluminal vesicles; ILVs)^23,24^. The direct fusion of mature MVBs with lysosomes that contain lipases, lysozyme and other hydrolytic enzymes facilitates the degradation of cargo, potentially terminating deleterious signal transduction by cell surface receptors such as epidermal growth factor receptor (EGFR)^16,25^. MVBs can also be driven towards the plasma membrane to release specific contents as exosomes^26,27^, an essential mode of intra-cellular communication^28–30^. Endocytic vesicular transport (EVT) pathway is known to use microtubules for motor- and Rab-protein regulated movements of endocytic vesicles containing cell surface receptor cargo^31–37^.

In this study, we identified a novel positive regulatory role for CPAP (centrosomal P4.1 associated protein; expressed by CENPJ gene)/SAS-4, a microtubule/tubulin binding essential centriole biogenesis protein^38^ in EVT of internalized epidermal growth factor receptor (EGFR). The centrosome, which consists of a pair of centrioles suspended in a peri-centriolar material (PCM), is the major microtubule organizing centers (MTOC) of mammalian cells^39^. Several centriole associated proteins can directly interact with microtubules^40–42^. Among them, CPAP is known to regulate centriolar microtubule growth to produce centrioles of optimum dimension^43^. Importantly, cytoplasmic microtubule function is critical for EVT and targeting of internalized cell surface receptors to lysosome^32,33,35^. To date, CPAP function is primarily attributed to centriole duplication, specifically restricting the centriole length, and ciliogenesis^38,44–47^. Here, by employing gain- and loss-of-function approaches as well as the ligand-bound EGFR intracellular-trafficking model, we assessed the role of CPAP in EVT and lysosome targeting of internalized cell surface receptor cargo. We found that CPAP is required for efficient transport of internalized EGFR to MVB and lysosome for degradation, and acts as a positive regulator of EVT. These novel findings add a new dimension to the cellular functions of a critical centriole biogenesis protein CPAP.

## Results

### CPAP overexpression causes the formation of punctate structures and increases the abundance of MVBs

The role of CPAP in centriole biogenesis and centrosome function is well established^38,41,48–52^. During our studies to determine the interaction between CPAP and other centriolar proteins, we observed that both HEK293T and HeLa cells transfected with GFP-CPAP **(Fig. 1 and Supplemental Fig. 1)** formed punctate vesicular structures spontaneously. These punctate structures were observed not only in most cells that were transiently transfected with GFP-CPAP (**Fig. 1A**), but also in cells that were stably expressing GFP-CPAP under a doxycycline (doxy)-inducible promoter upon doxy treatment (**Fig. 1B**). Immunofluorescence (IF) microscopy of both HEK293T and HeLa cells that transiently express myc-tagged CPAP (myc-CPAP) also revealed vesicular structures but with a distinct membrane-structure localization of the exogenously expressed CPAP **(Figs. 1C and 1D).** These observations indicate that CPAP overexpression associated formation of puncta may not be mere protein aggregates formed due to higher expression levels of GFP- and myc-tagged CPAP and, perhaps, CPAP localizes to intracellular vesicular structures.

**Fig. 1:**
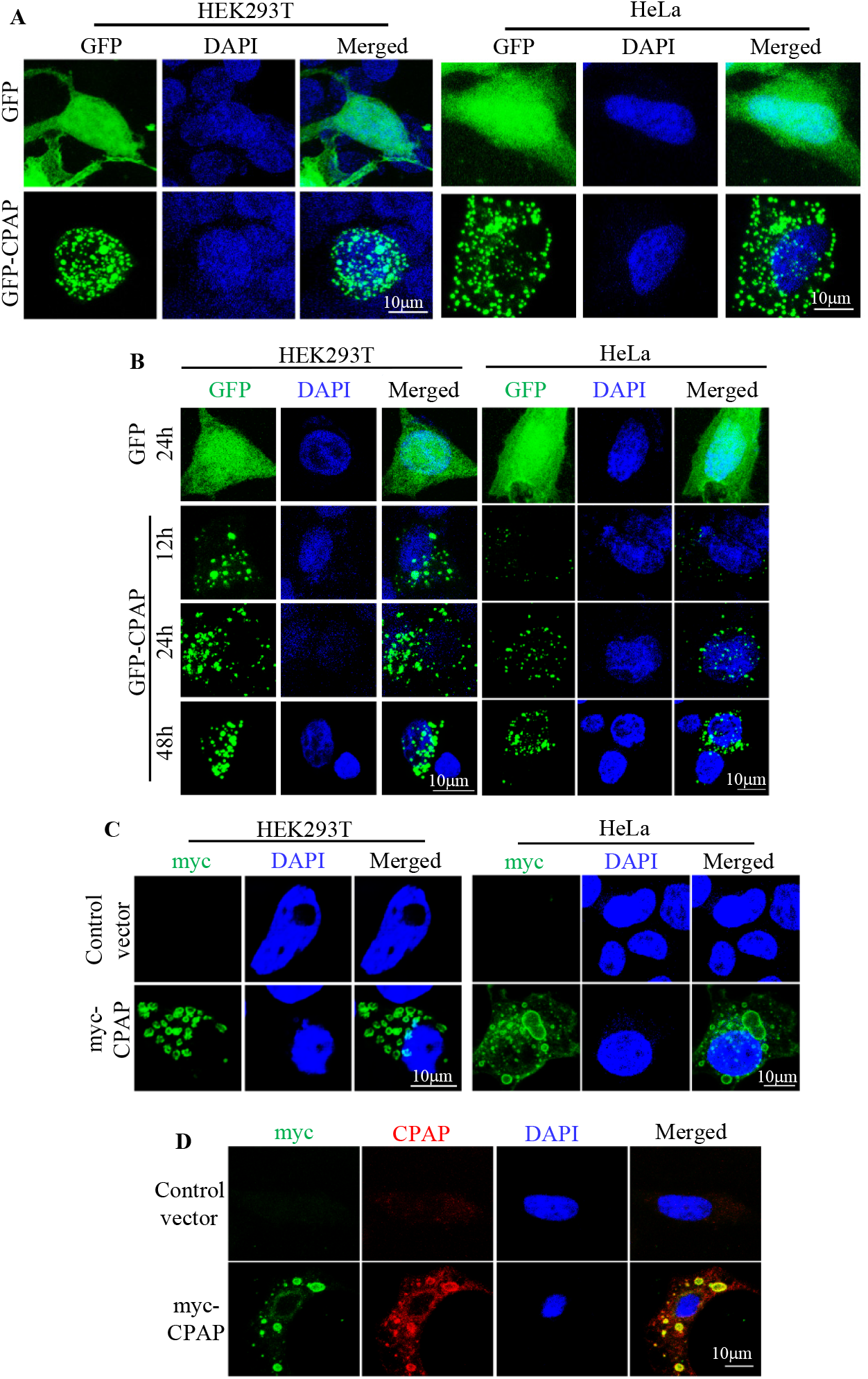
CPAP overexpression causes the formation of vesicular structures. **A.** HEK293T and HeLa cells were transiently transfected with GFP or GFP-CPAP expression vectors for 24h and imaged using Zeiss 880 confocal microscope. **B.** HEK293T and HeLa cells expressing GFP or GFP-CPAP under doxycycline (doxy) -inducible promoter were imaged. **C.** HEK293T and HeLa cells were transfected with control or myc-CPAP expression vectors for 24h and subjected to staining using anti-myc antibody and imaging. **D.** HeLa cells were transfected with control or myc-CPAP expression vectors for 24h and subjected to staining using anti-myc and anti-CPAP antibodies and imaging. Supplemental Fig. 1 shows IB analysis of various cells for GFP-CPAP and myc-CPAP expression.

### CPAP overexpression causes increase in the abundance of MVB-like structures

Although CPAP overexpressing cells that were used in previous reports appeared to show similar puncta formation as shown in Fig.1 ^42,44,45^, the significance of these structures were not discussed or further investigated. We have performed transmission electron microscopy (TEM) to assess the morphological features of CPAP-overexpressing cells. As shown in **Fig. 2A**, HEK293T cells transfected with GFP-CPAP and myc-CPAP showed the presence of large, electron dense bodies. These electron dense structures have the characteristics of MVB, which are multiple smaller vesicles called intraluminal vesicles (ILVs) bound by a limiting membrane^17,18,25,53^. Interestingly, CPAP overexpression associated MVB-like structures were found to be not only higher in numbers, but also larger in dimensions, than the ones observed in control cells. These observations prompted us to stain GFP-CPAP expressing cells with markers of EE (EEA1), MVB/ LE (CD63) and endolysosome/lysosomes (LAMP1) to assess if the exogenously expressed CPAP localizes to these vesicles of endocytic origin. Maximum projection and single Z scan images show that a considerable number of GFP+ vesicles partially overlap with EEA1+ and CD63+ vesicular structures **(Fig. 2B)**. However, very few GFP+ vesicles showed such overlap or colocalization with LAMP1+ vesicles. Overall, these results suggest, irrespective of the localization of CPAP to specific functional stages of endosomes, higher CPAP level contributes to the formation of MVB-like structures.

**Fig. 2:**
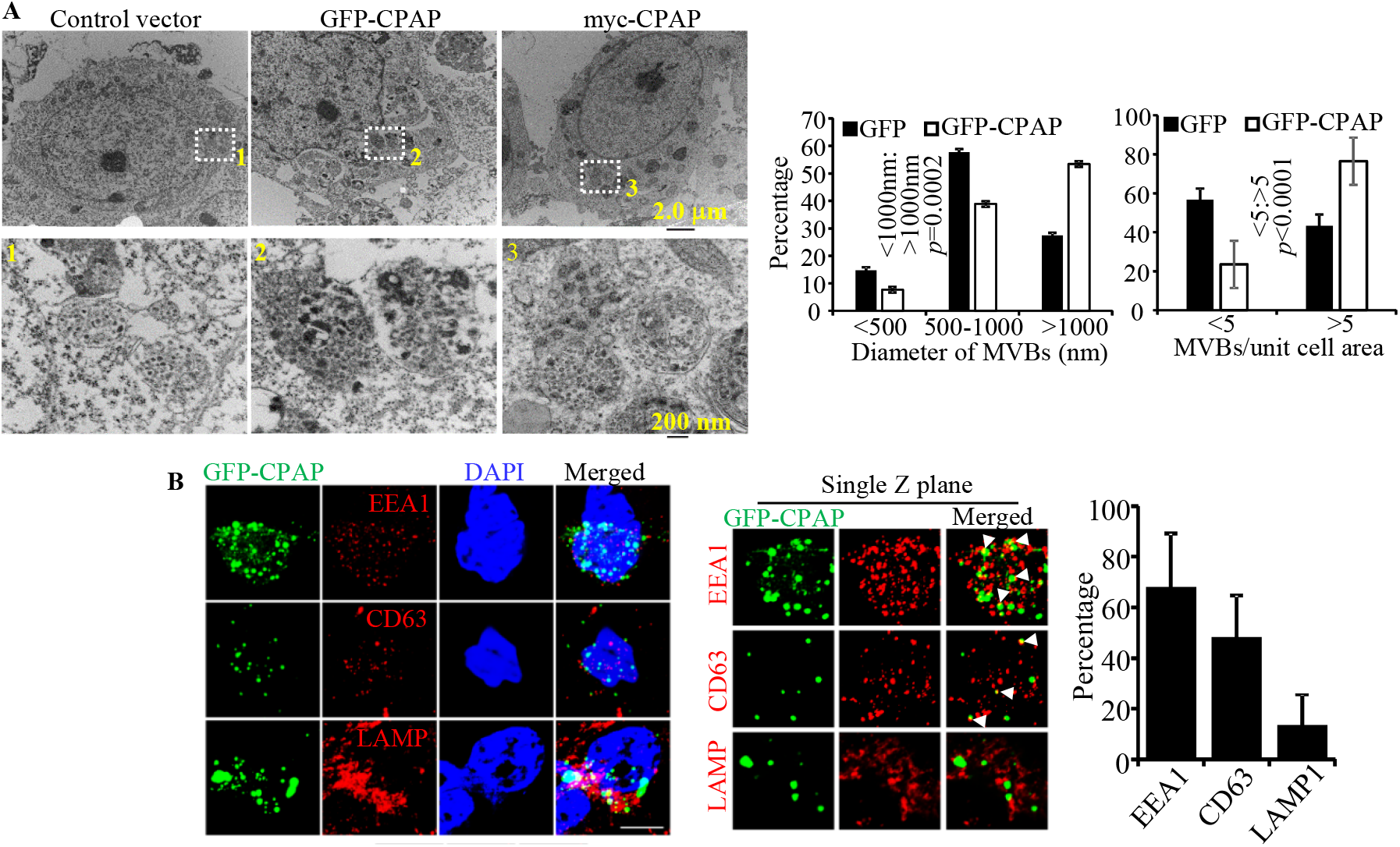
CPAP overexpression causes the formation of MVB-like structures. **A.** HEK293T cells were transfected with control plasmid, or GFP-CPAP or myc-CPAP expression vectors for 24h as described for Fig. 1, fixed with paraformaldehyde and processed for TEM, and representative images are shown in upper panels. Numbers of MVB-like structures per unit cell [GFP n=13; GFP-CPAP n=43] and diameters of individual MVB like structures [GFP n=24; GFP-CPAP n=161] were calculated from three separate experiments for control and GFP-CPAP vector transfected cells and the Mean ± SD values (lower panels) are shown. *p*-value is by chi2 test comparing the ratios of MVBs with <1000nm and >1000nm sizes or ratios of cells/images with <5 and >5 MVBs between control and GFP-CPAP cells. Note: MVB number and size were determined in transfected cell preparations containing a mixture of cells with and without GFP expression in an unbiased manner for both groups. **B**. HEK293T cells were transfected with GFP-CPAP expression vector for 24 h and subjected to staining for EEA1 (early endosomes), CD63 (MVBs), and LAMP1 (lysosomes and late endosome), and images were acquired using Airyscan unit of Zeiss 880 microscope. Maximum projection images of Z-stacks (left panel) and single Z planes of relevant images (middle panel), and quantitation of percentage of GFP+ puncta associated/co-localized with EEA1, CD63 or LAMP positive vesicles in at least 25 cells/group (right panel) are shown.

### Cell surface receptor EGFR is detected in CPAP overexpression induced puncta

If CPAP overexpression causes the formation of MVBs and, potentially, constitutive activation of EVT, then cell surface receptors such as EGFR could be continuously targeted to lysosome for degradation in CPAP overexpressing cells. To test this notion, first we examined if EGFR can be found in CPAP overexpression-induced vesicular structures of cells that are grown in complete medium, which contains low levels of EGF of FBS origin. **Fig. 3A**, single Z-plane images particularly, show that endogenous EGFR is detectable within or associated with several of the CPAP overexpression-induced spontaneous vesicular structures in both HEK293T and HeLa cells. We then examined if ligand activation induced EGFR (internalized Alexa-fluor 555-linked EGF) localizes to CPAP-overexpression induced vesicular structures. As observed in **Fig. 3B**, considerable amount of internalized, fluorescent EGF-bound EGFR containing vesicular structures were associated with GFP-CPAP+ puncta. This suggests that vesicular transport and lysosomal degradation of EGFR is more rapid in CPAP overexpressing cells compared to control cells. Hence, we examined the cellular levels of EGFR in doxy-treated HeLa cells that are stably expressing GFP-CPAP under a doxy-inducible promoter. As observed in **Fig. 3C,** cellular levels of EGFR, detected by IB, were diminished considerably upon induction of CPAP expression by doxy, in a dose-dependent manner. In addition, FACS analysis revealed lower levels of EGFR on the cell surface **(Fig. 3D)**. These results suggest that, by facilitating EVT function, CPAP may have a positive regulatory role in the homeostasis of EGFR-like cell surface receptors.

**Fig. 3:**
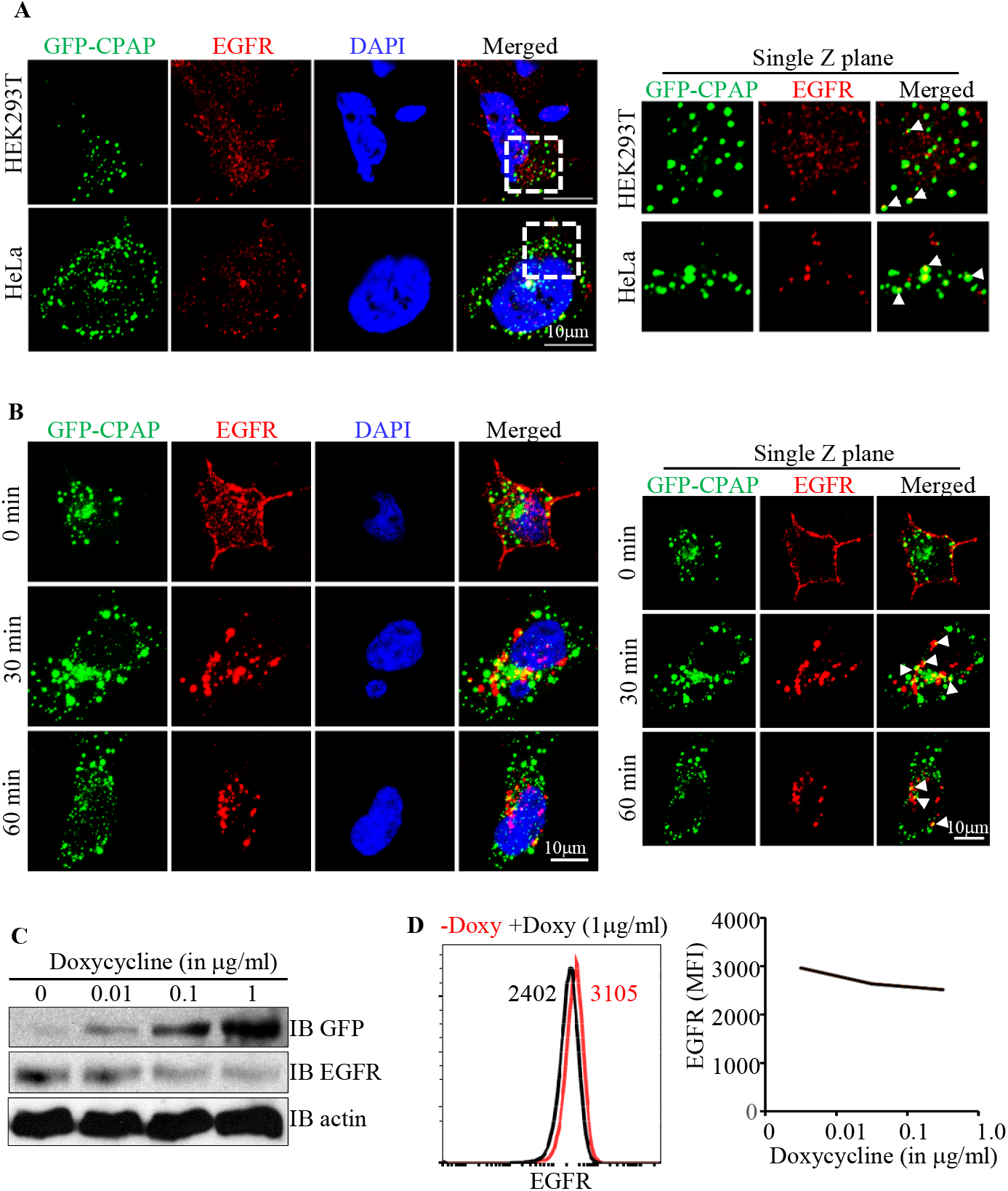
EGFR is detected in CPAP overexpression associated vesicular structures and higher levels of CPAP diminish the cellular levels of EGFR. **A.** HEK293T and HeLa cells expressing GFP-CPAP under doxy-inducible promoter were stained using anti-EGFR antibody, 24 h after induction of protein expression with doxy (0.1 μg/ml). Maximum Z-projection images (left panel) and single Z plane of relevant images (right panel) are shown. White arrows show examples of colocalization. **B.** GFP-CPAP expressing HeLa cells were treated with Alexa fluor 555-conjugated EGF for different time-points and subjected to confocal microscopy to detect ligand-bound EGFR. Maximum Z-projection (left panel) and single Z-plane (right panel) images of representative cells are shown. **C.** HeLa cells stably expressing GFP-CPAP under doxy-inducible promoter were left untreated or treated with varying amounts of doxy for 24 h and subjected to WB to detect GFP-CPAP, EGFR and β-actin. **D.** Untreated and Doxy treated cells were surface stained using fluorchrome linked anti-EGFR antibody and examined by FACS. Representative overlay graph with MFI values (left panel) and mean MFI values of cells (3 independent experiments) exposed to different doses of Doxy for 24 h (right panel) are shown. Original scanned x-ray film images or ChemiDoc imager acquisition images of relevant IB panels are included as supplemental figure 6.

### CPAP depletion causes increased cellular levels of total EGFR

Since CPAP overexpression diminishes EGFR levels, we studied if CPAP depletion impacts the surface and cellular levels as well as the internalization and degradation dynamics of EGFR. First, HeLa cells stably expressing control (scrambled) shRNA and CPAP-shRNA (CPAP-depleted) were examined for cellular **(Fig. 4A)** and surface **(Fig. 4B)** levels of EGFR by IB and FACS. Relative to control shRNA expressing cells, the cellular and surface levels of this receptor were higher in CPAP-depleted cells. Next, these cells were treated with the ligand, EGF for different durations, under cycloheximide (chx) treatment to block nascent protein synthesis, and the surface and cellular levels of EGFR were determined. **Fig. 4C** shows that the cell surface levels of EGFR are relatively higher in CPAP-depleted cells compared to control cells at all time-points. However, the surface levels of EGFR in both control and CPAP-depleted cells showed similar degrees of decrease after EGF treatment, as indicated by the fold change in overall mean fluorescence intensity (MFI). This suggests that, although the basal levels of EGFR are lower in CPAP-depleted cells, the rate of receptor internalization is not significantly altered upon CPAP depletion. IB analysis of EGF treated cells revealed higher cellular levels of EGFR in CPAP depleted cells irrespective of the time-point tested (**Fig. 4D**), suggesting that EGFR degradation pathway is negatively impacted by CPAP depletion. Notably, while majority of the cellular EGFR appears to have degraded in control cells within 120 min post-treatment compared to 0 min time point, CPAP-depleted cells showed the persistence of significant amounts of EGFR at this time-point suggesting its diminished degradation kinetics when CPAP levels are low. To determine if altered EGFR levels under CPAP depletion translate to the degree of its function, cellular levels of phosphorylated forms of EGFR and downstream target AKT were determined. As shown in **Fig. 4E**, levels of phospho-EGFR and phospho-AKT were higher in CPAP-depleted cells, upon EGF treatment, compared to control cells. To rule out the possibility of non-specific effects due to viral vector mediated shRNA expression and puromycin selection, similar experiment was conducted after transient depletion of CPAP expression using siRNA which targets a different region. As observed in **Fig. 4F and 4G**, similar to shRNA expressing cells, HeLa cells treated with CPAP-specific siRNA showed considerably higher basal cellular and surface levels of EGFR. We, then treated HeLa cells with CPAP-specific siRNA and transfected with siRNA-resistant GFP-CPAP cDNA vector^44^ to assess if reintroduction of CPAP function restores EGFR levels. These cells were treated with EGF as done for Fig. 4D prior to detection of cellular EGFR levels by IB. Transient depletion of CPAP using siRNA, similar to its stable depletion (Fig. 4D), resulted in higher cellular levels of EGFR, compared to control cells **(Fig. 4H)**. CPAP siRNA treated cells also showed diminished ligand engagement-induced degradation of this receptor, as indicated by little or no reduction in cellular levels after EGF treatment. Importantly, reintroduction of CPAP levels appears to restore the ability of ligand treated cells to degrade EGFR as suggested by its diminished cellular levels, post-ligand engagement as compared to 0 h time-point. Overall, these observations show defective degradation of internalized EGFR, and its sustained/prolonged function, when CPAP levels are low. These results suggest that CPAP positively contributes to the homeostasis of EGFR and, potentially, other similar cell surface receptors. In fact, **Supplemental Fig. 2**, shows that CPAP depletion also impacts the cellular levels of TGF-β receptors, which are also known to be targeted to the lysosome for degradation^54^.

**Fig. 4:**
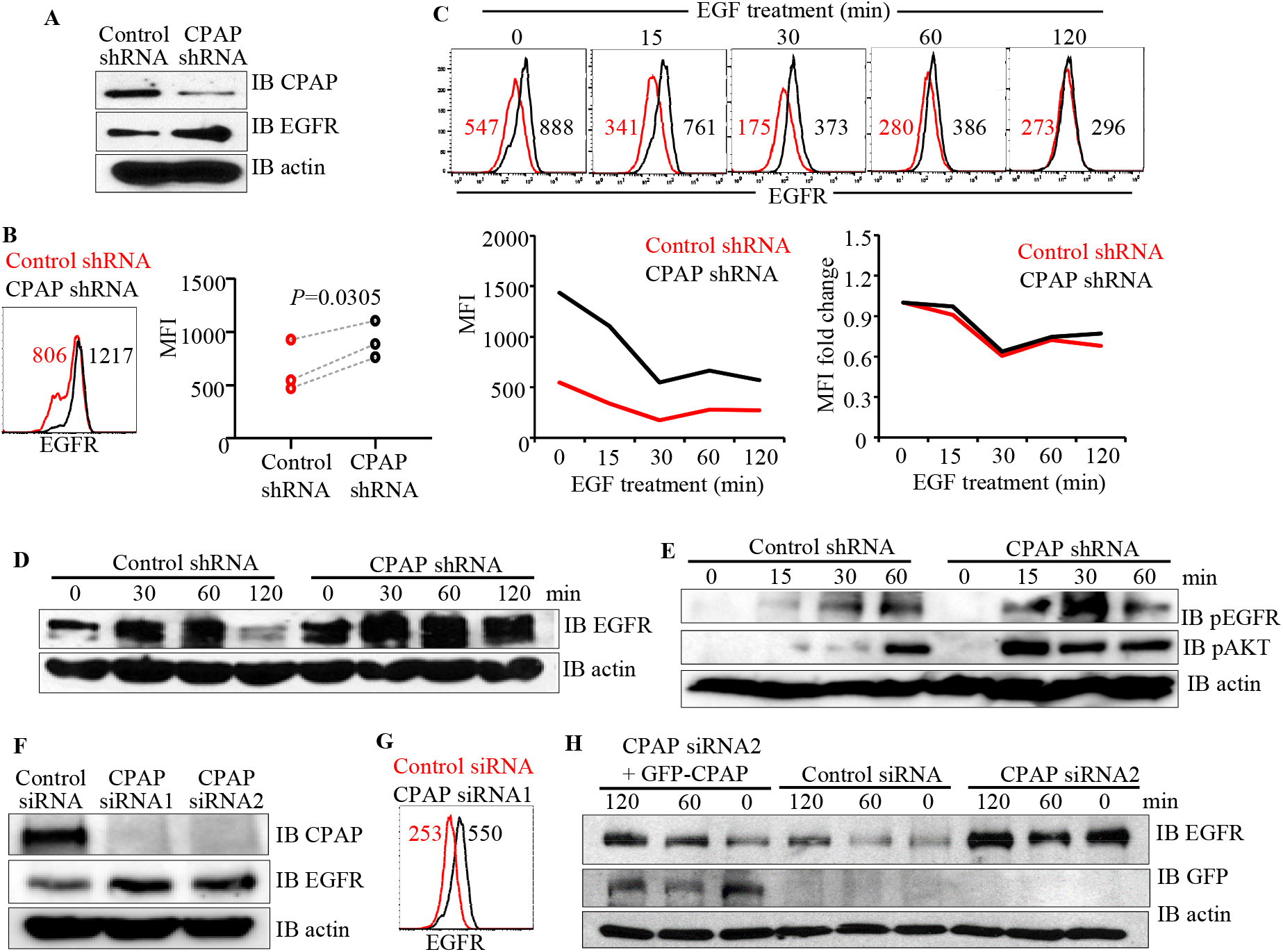
CPAP depletion increases surface and cellular levels of EGFR. HeLa cells stably expressing control-shRNA or CPAP-shRNA were examined for CPAP depletion and EGFR levels by WB (**A**) and surface EGFR levels by FACS (**B**). Panel B shows representative overlay graph (left panel) and MFI values from 3 independent experiments. *P*-value by paired t-test. Cells were subjected to serum starvation overnight, treated with cycloheximide for 1h and EGF, washed and incubated for indicated durations, and subjected to FACS analysis to detect surface levels of EGFR after staining using anti-EGFR antibody. Representative overlay graphs for each time-point (**C, upper panel**), mean MFI values of three independent assays (**C, lower left panel**) and EGF treatment induced fold changes in EGFR specific MFI values, relative to 0 min time-point, (**C, lower right panel**) are shown. **D.** Cells stimulated using EGF as described in panel C were subjected to WB to detect EGFR and β-actin. **E.** Cells stimulated using EGF for different durations as described in panel C and subjected to WB to detect phospho-EGFR, phospho-AKT and β-actin. **F.** HeLa cells were treated with control-siRNA, CPAP-siRNA1 or CPAP-siRNA2 for 72h and subjected to IB to detect cellular CPAP, EGFR and β-actin. **G.** siRNA treated cells were subjected to FACS to detect surface levels of EGFR and representative overlay graph with MFI values is shown. **H.** siRNA treated cells that exogenously express siRNA-resistant GFP-CPAP under doxy-inducible promoter, were treated with EGF as described for panel D and subjected to IB analysis to detect GFP-CPAP, EGFR and β-actin. GFP-CPAP expressing cells were treated with doxy for 24h before initiating this assay. Original scanned x-ray film images or ChemiDoc imager acquisition images of relevant IB panels are included as supplemental figure 6.

### Targeting of cell surface receptors to EE is not affected in CPAP depleted cells

Internalized ligand activated receptors are routed to EE, following which they are targeted to the lysosome via MVB for degradation or recycled back to the surface^55^. CPAP-depleted cells do not appear to be defective in receptor internalization (Fig. 4). Hence, we examined if the earliest events of EVT such as EE targeting of ligand bound EGFR, most of which is known to be targeted to lysosome, were affected by CPAP depletion. Control and CPAP depleted HeLa cells that were treated with Alexa flour 555-linked EGF were stained for EEA1 and subjected to confocal IF microscopy to determine the early endosome localization of ligand bound EGFR. Maximum projection and single Z images, and quantitation of ligand-bound EGFR localization to EEA1+ vesicles showed that EGFR localizes to EEA1 positive vesicular structures at a comparable degree in both control and CPAP depleted cells **(Fig. 5A)**. This suggests that CPAP does not have a regulatory effect on internalization or transport of receptors to EE. We also examined if the localization of internalized ligand (transferrin; Tfn)-bound transferrin receptor (TfnR), which is primarily recycled back to the cell surface and does not enter the degradation pathway^55^, to the EE is affected by CPAP depletion. Control and CPAP depleted cells were treated with Tfn for different durations and subjected to confocal microscopy after staining for TfnR and EEA1. **Fig. 5B** shows that, similar to EGFR, TfnR is routed to the EE, as indicated by colocalization with EEA1, at comparable degrees in control and CPAP depleted HeLa cells. Further, FACS analysis showed that, unlike the EGFR (Fig. 4), both control and CPAP depleted HeLa cells maintain comparable levels of surface TfnR under steady state and ligand treatment (**Supplemental Fig. 3**) suggesting that this receptor is recycled back to the cell surface at comparable rates in control and CPAP depleted cells. Overall, in conjunction with the results of Figs. 4 showing the cellular levels and subcellular dynamics of EGFR, these observations suggest that CPAP, perhaps, has a regulatory role in the transport of internalized cell surface receptor cargo from EE to late endosome/MVB and lysosome, but not to the EE or recycling back to the surface.

**Fig. 5:**
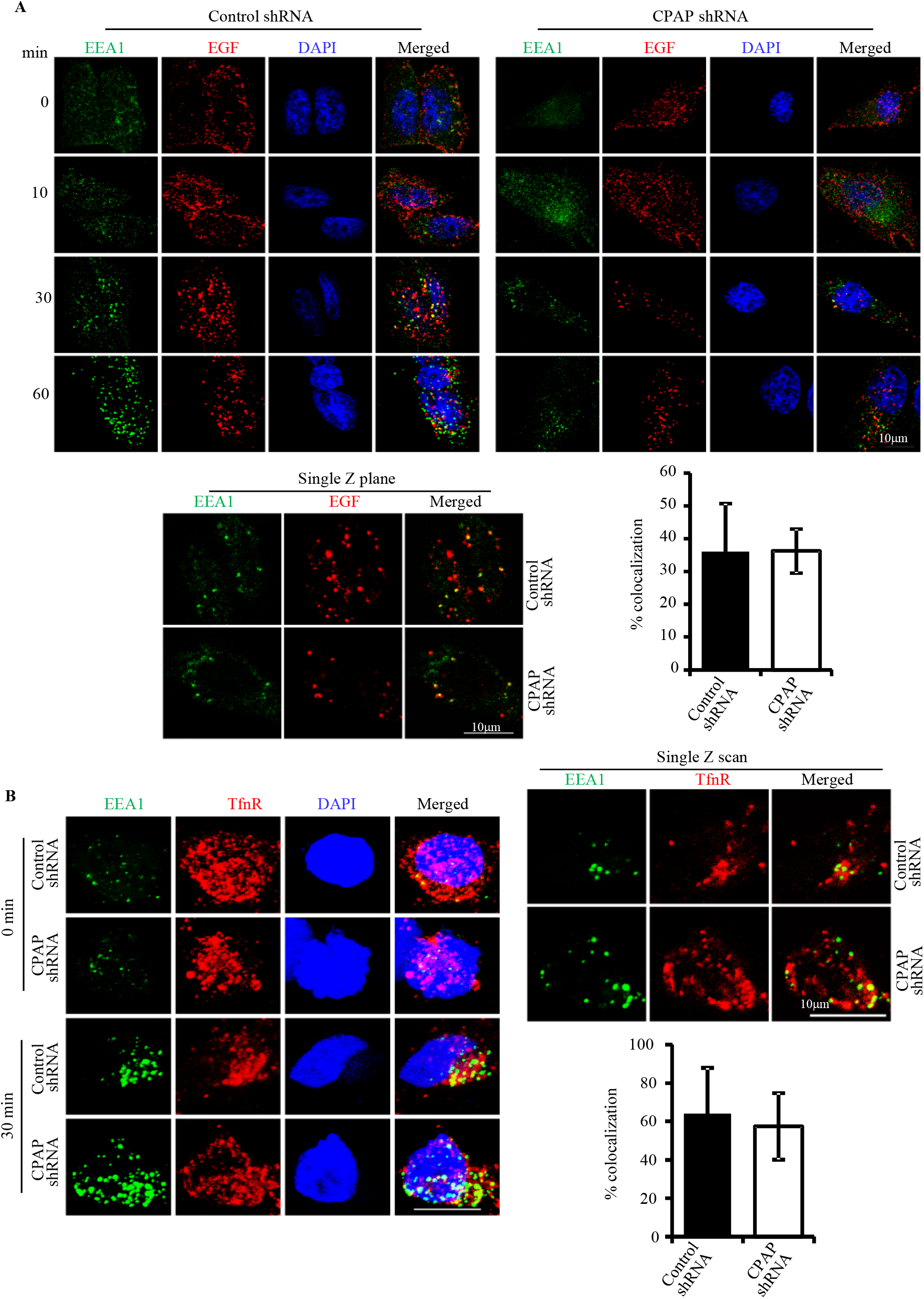
CPAP depletion has no impact on trafficking of cell surface receptor to early endosomes. **A.** HeLa cells expressing control-shRNA or CPAP-shRNA were incubated with Alexa fluor 555-conjugated EGF ligand and left on ice for 1h. Cells were washed with serum free media and transferred to 37°C to initiate receptor internalization. Cells were fixed at indicated timepoints, permeabilized and stained for EEA1 to mark early endosomes. Images were acquired as Z-stacks using Zeiss 880 microscope. Maximum projection images of cell harvested at different time-points (left panel), and single Z plane of relevant images (upper right panel) and mean ± SD values (30 min time-point) of percentage EGF+ vesicular structures that co-localize with EEA1 staining (lower right panel) from three independent experiments are shown. Vesicular structures of a single Z-plane/cell (at least 15 cells/experiment) were counted. **B.** HeLa cells expressing control-shRNA or CPAP-shRNA were incubated with holotransferrrin and processed similarly to that for panel A and stained for TfR and EEA1. Maximum projection images of cells harvested at different timepoints (left panel), and single Z plane of relevant images (upper right panel) and mean ± SD values (30 min time-point) of percentage TfR+ vesicular structures that co-localize with EEA1 staining (lower right panel) from three independent experiments are also shown. Vesicular structures of a single Z-plane/cell (at least 15 cells/experiment) were counted.

### CPAP depletion results in defective trafficking of EGFR to lysosomes

It is well established that ligand engaged cell surface receptors such as EGFR are degraded in the lysosome^16^. Since our results showed diminished degradation and higher cellular and surface levels of EGFR and unaltered EE targeting of internalized receptors in CPAP depleted cells, we examined if targeting of ligand engaged EGFR to the lysosomes is affected in these cells. HeLa cells stably expressing control-shRNA and CPAP-shRNA were treated with Alexa fluor 555-linked EGF on ice, washed and incubated at 37°C for different time-points and stained for the endolysosome/lysosome marker LAMP1, and subjected to confocal microscopy. As observed in maximum projection images of **Fig. 6**, control cells showed profound perinuclear clustering of ligand-bound EGFR and colocalization of this receptor with LAMP1, particularly at 60 min post-ligand treatment. However, compared to these control cells, internalized EGFR in CPAP depleted cells showed reduced perinuclear clustering and colocalization with LAMP1. Single Z scan images and quantification of ligand-bound EGFR-containing structures localized to LAMP1+ vesicles at 60 min time point clearly show inefficient trafficking of EGFR to the lysosomes in CPAP-depleted cells. Of note, lysosome targeting of internalized EGFR appears to be delayed in CPAP depleted cells as indicated by relatively higher perinuclear clustering of EGFR in these cells at later time-point (120 min) compared to the 60 min time-point. Experiments using cells that are depleted of CPAP using siRNA also showed similar defective targeting of EGFR to the lysosome **(Supplemental Fig. 4)**. Overall, these observations suggest that defective transport of internalized EGFR from EE to lysosomes is the reason for higher total- and phospho-EGFR levels in cells under CPAP deficiency, and CPAP positively regulates lysosome targeting of internalized protein cargo for degradation.

**Fig. 6:**
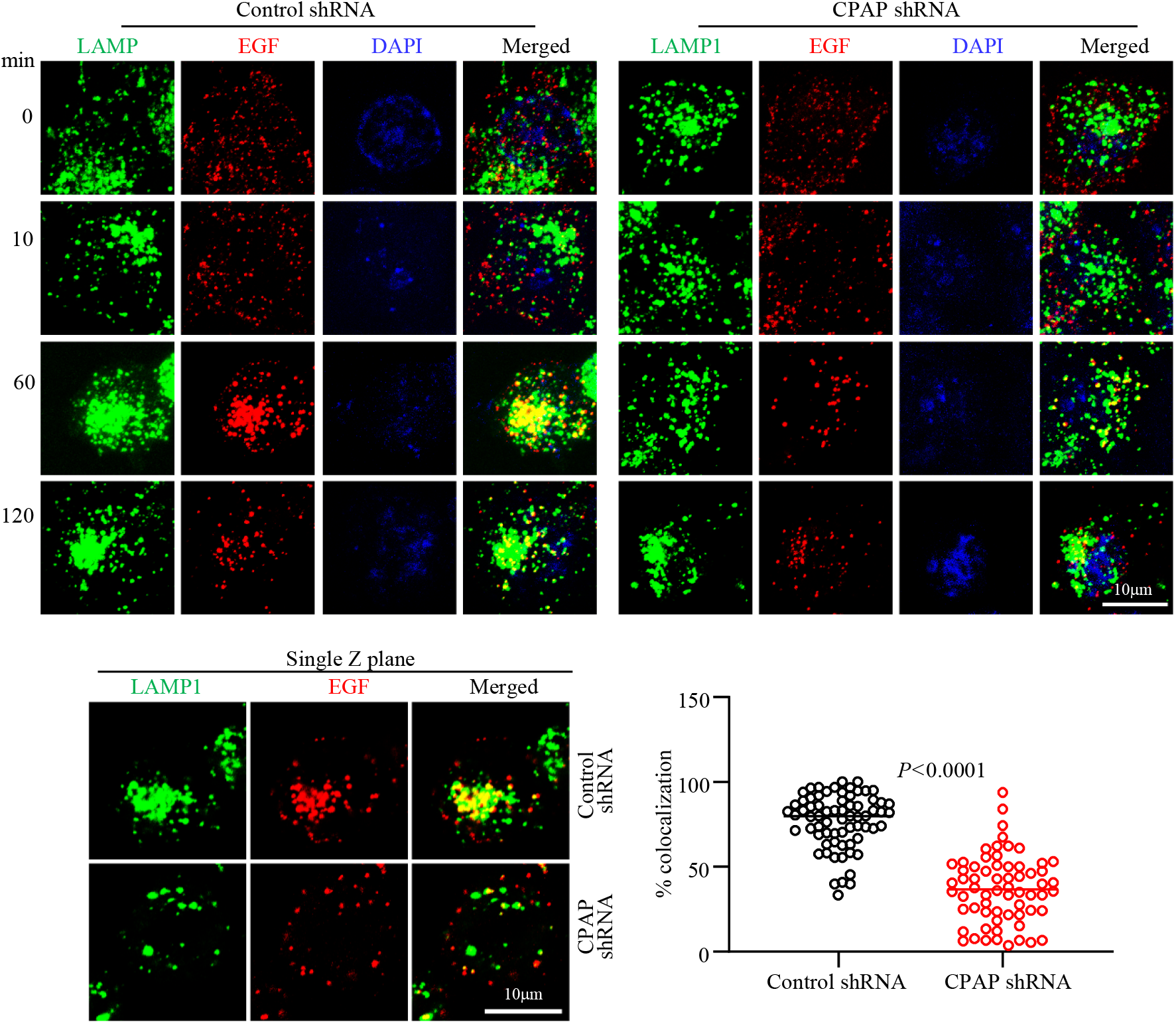
CPAP depletion results in defective targeting of internalized cell surface receptor to the lysosome. HeLa cells expressing control-shRNA or CPAP-shRNA were incubated with Alexa fluor 555-conjugated EGF ligand and left on ice for 1h. Cells were washed with serum free media and transferred to 37°C to initiate receptor internalization. Cells were fixed at indicated timepoints, permeabilized and stained for LAMP1 to mark lysosomes, and images were acquired as Z-stacks using Zeiss 880 microscope. Maximum projection images of cells harvested at different time-points (left panel), and single Z-plane of relevant images (upper right panel) and percentage of EGF+ vesicular structures that co-localize with LAMP1+ staining at 60 min time-point (lower right panel) are shown. Vesicular structures of a single Z-plane/cell (at least 20 cells/experiment) were counted.

### CPAP depletion results in defective trafficking of EGFR to MVB/late endosomes

Late endosomes/MVBs, which are matured from EE, fuse with lysosomes to form endolysosomes for delivery of internalized cargo proteins for their degradation^56^. Since we observed: 1) higher abundance of MVB in CPAP overexpressing cells (Fig. 2A) and 2) lysosome targeting of EGFR is negatively impacted by CPAP deficiency, we examined if the transport of internalized EGFR from EE to LE/MVB is impacted by CPAP levels. HeLa cells stably expressing control-shRNA and CPAP-shRNA were treated with Alexa fluor 555-linked EGF for different durations and stained for MVB/late endosome marker CD63 for confocal microscopy. Similar to LAMP staining, maximum projection images of CD63 staining **(Fig. 7A)** showed profound perinuclear clustering of ligand-bound EGFR and localization of this receptor in CD63+ vesicular structures in control cells, particularly at 60 min post-treatment. However, in CPAP depleted cells, profoundly less perinuclear clustering and colocalization of ligand-bound EGFR with CD63 was observed at this time point. Single Z scan images and the quantification of ligand-bound EGFR-containing structures localized to CD63+ vesicles at 60 min time-point confirms significantly diminished transport of ligand bound EGFR to MVBs in CPAP-depleted cells. Similar observations were made with CPAP specific siRNA treated cells that were examined for the localization of ligand bound EGFR to CD63+ vesicles **(Supplemental Fig. 5)**. Overall, these results suggest that CPAP facilitates targeting of the ligand bound EGFR cargo from EE to LE and lysosome, perhaps by affecting MVB formation. In fact, TEM analysis revealed significantly lower number of electron dense and MVB-like structures, endolysosome and lysosomes in CPAP depleted cells compared to control cells upon EGF treatment **(Fig. 7B)**. These observations suggest diminished ability to form MVBs in CPAP depleted cells is responsible for the defective transport of EGFR from EE to lysosome and its degradation.

**Fig. 7:**
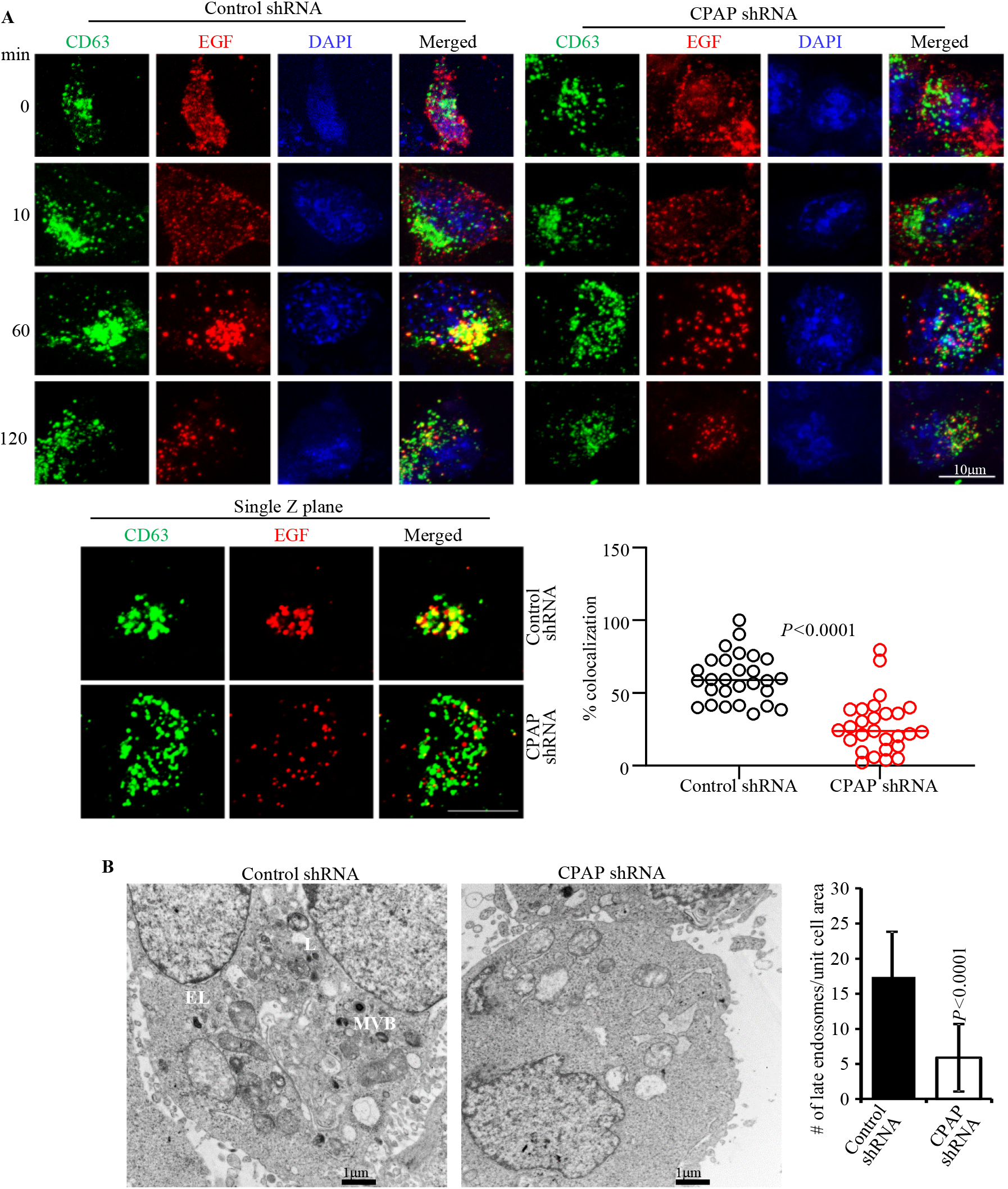
CPAP depletion results in defective trafficking of internalized cell surface receptor to MVB/late endosome. **A.** HeLa cells expressing control-shRNA or CPAP-shRNA were incubated with Alexa fluor 555-conjugated EGF ligand and left on ice for 1h. Cells were washed with serum free media and transferred to 37°C to initiate receptor internalization. Cells were fixed at indicated time-points, permeabilized and stained for CD63 to mark MVB/late endosome. Images were acquired as Z-stacks using Zeiss 880 microscope. Maximum projection images of cells harvested at different time-points (left panel), and single Z-plane of relevant images (upper right panel) and percentage of EGF+ vesicular structures that co-localize with CD63+ staining at 60 min time-point (lower right panel) are shown. Vesicular structures of a single Z-plane/cell (at least 25 cells/group) were counted. **B.** HeLa cells expressing control-shRNA or CPAP-shRNA were treated with EGF for 1 h and subjected to TEM analysis, and representative images (left panel) and mean ± SD values (right panel) of number of late endosome [MVBs (M) and endo-/lysosomes (electron dense bodies; EL and L) are shown. Vesicles were quantified for at least 15 cells/group.

## Discussion

Here, we establish a novel regulatory role for CPAP, an essential centriole biogenesis protein^38^, in endosome maturation and lysosome targeted EVT of ligand engaged cell surface receptor EGFR. CPAP is a microtubule and α-tubulin binding protein and its tubulin-binding function^52,57,58^ is important for its function on the centrioles, especially in restricting the centriole length to <500nm^44–46,58^. Overexpression of CPAP can cause aberrant long centrioles of up to 3 μm^44–46^. In addition, CPAP is required for spindle orientation, which defines normal and asymmetric cell divisions^59^. Mutations in the human CENPJ gene have been associated with microcephaly and Seckel syndrome^60–62^. A hypomorphic mouse mutant of the CENPJ gene not only developed microcephaly of the brain but also accumulated centriole duplication defects^63^. Recently, it was also shown that CPAP regulates progenitor divisions and neuronal migration in the cerebral cortex. Centrioles can serve as basal bodies, which template the formation of cilia^64^. More recently, CPAP has been shown to regulate the length of cilia^44,46,47^, perhaps through its centriolar function. Expression of CPAP is cell-cycle regulated^45^ and it also determines the timing of cilium disassembly^65^. In this study, however, we have uncovered a novel function of this centriole biogenesis protein as a positive regulator of the EVT of EGFR-like cell surface receptors and their homeostasis.

MVBs serve as critical intermediates towards curbing receptor-mediated signal transduction pathways by sequestering the cargo of EE into MVBs and routing these proteins for lysosomal degradation^25^. Our results show that CPAP does not regulate the internalization or routing of ligand-engaged EGFR into EE. Further, routing for cell surface receptors such as TfnR that are recycled back to the cell membrane from the early endosome also does not appear to be regulated by CPAP. However, our results demonstrate that MVB formation and the transport of receptors which are destined for lysosome degradation, is facilitated by CPAP. This notion has been substantiated by our observation that ligand engaged EGFR colocalization with CD63+ and LAMP+ vesicular structures in cells is profoundly diminished when CPAP is depleted. Perinuclear clustering is the key feature of LE and lysosome entry of protein cargo that is targeted for lysosomal degradation^66^. Hence the notion that CPAP facilitates endosome maturation is further supported by our microscopy studies revealing defective perinuclear clustering of EGF+ vesicles in CPAP-depleted cells as compared to control cells. Our ultra-structural studies revealing that higher CPAP expression causes increase in the number and size of MVB-like structures and CPAP depletion results in diminished LE/MVB and endolysosome structures suggests that in fact, fewer endocytic vesicles are generated when CPAP expression is low, leading to a blockade in the sequential progression of cargo transport from EE to LE, and to the lysosome.

While previous reports were primarily focused on the centriolar localization and function of CPAP, considerable amount of CPAP is also found in the cytoplasmic pool^59^. In fact, cytoplasmic-centrosomal shuttling of this protein has been described as a mechanism of regulating centriole elongation^45,59^. However, a role of this protein in endosome maturation has never been described. Our studies not only show that overexpressed CPAP is found on the vesicular structures, but also the CPAP protein is detectable with in endocytic vesicles. We also found that internalized EGFR is associated with CPAP overexpression induced puncta. However, if the endosome localization of CPAP is required for the vesicular transport function or MVB formation needs to be investigated. One potential molecular mechanism could be that CPAP is, directly or indirectly through interaction with EVT associated proteins, involved in MVB biogenesis and facilitates the lysosomal targeting of the receptor cargo. Other possible mechanism could be through Rab proteins. Rab proteins regulates several steps of the vesicle trafficking processes to release the cargo to be sorted for degradation^67^. It has been shown that Rab5 dynamically fluctuates on individual EE, linked by vesicle fusion and fission events along with the degradative cargo such as EGFR and concentrates in progressively fewer and larger endosomes^68,69^. This process occurs when endosomes migrate from the cell periphery to the perinuclear region along the MT where Rab5 is rapidly replaced by Rab7. Since CPAP is a microtubule-associated protein, with high affinity to bind tubulin, it is possible that this function is compromised when CPAP levels are low resulting in a collapse in the endocytic vesicle tethering function, defective cargo movement and vesicle maturation along the microtubule filaments. These aspects need to be addressed in future studies.

In conclusion, our study identifies a novel function for a quintessential centriole biogenesis protein in endosome maturation during the vesicular transport of cell surface receptor cargo targeted for degradation. A wide repertoire of centrosome-associated cellular processes including centriole duplication, mitotic progression, spindle orientation, and ciliogenesis have been attributed to CPAP. However, whether this newly identified role of CPAP in endosome maturation is required for these processes or is an independent function remains to be studied. Further, whether the centriole-localization of CPAP and its tubulin-binding ability attributes to this newly discovered function on vesicular transport remains to be determined. Nevertheless, novel observations described here will pave the way for new studies to determine if and how the fundamental cellular processes such as centriole duplication and endosome maturation are coupled through CPAP. Importantly, these observations could, potentially, also help better explain the molecular mechanisms of aberrant CPAP expression/function-associated microcephaly, mitotic and spindle positioning errors and ciliopathies.

## Materials and methods

### Cell lines

HEK293T (National Gene Vector Biorepository) and HeLa (ATCC) cells were used in this study. These cells were cultured in DMEM media supplemented with 10% FBS, sodium pyruvate, sodium bicarbonate, minimum essential amino acids and antibiotics. Transfection of plasmids was performed using calcium phosphate reagent or TransIT 2020 reagent from Mirus Bio LLC while siRNA was transfected using the TransIT siquest reagent from Mirus Bio LLC.

### Plasmids and reagents

GFP-CPAP and myc-CPAP cDNA expression vector^45^ used in this study were kindly provided by Dr. T.K. Tang, University of Taipei, Taiwan. For stable depletion of CPAP, validated lentiviral constructs (pLKO.1) expressing Mission shRNA targeting the following region 5’-GCTAGATTTACTAATGCCA-3’ in CPAP or scrambled shRNA were purchased from Sigma-Aldrich. For generation of stable cells, lentivirus was generated in 293T cells using accessory plasmids dR8.2 and VSV-G. Target cell line of interest was transduced with virus and selected for shRNA expression by treatment with the drug puromycin (2μg/ml). RNAi resistant construct for expressing GFP-CPAP was kindly provided by Dr. Pierre Gonczy, Swiss Institute for Experimental Cancer Research, Switzerland and the CPAP siRNA targeting sequence has been reported earlier^59^.Primary antibodies used in this study: anti-CPAP (Proteintech), -GFP (Santa Cruz Biotech) -actin (Proteintech), -EGFR (Santa Cruz Biotech), TfnR-Alexa fluor 647 (Biolegend) -CD63 (BD Biosciences), -EEA1 (Bethyl labs), -Rab5 (Santa Cruz Biotech), -Rab7 (Proteintech) and -EGFR-PE (Biolegend). Cycloheximide and doxycycline were purchased from Sigma-Aldrich. Unconjugated EGF ligand was purchased from Tonbo biosciences, Alexa fluor-555 conjugated EGF (Invitrogen), and holotransferrin ligand was from R&D Biosystems. siRNA targeting different regions of CPAP (CPAP-siRNA1: 5’-CCCAATGGAACTCGAAAGGAA-3’ or CPAP-siRNA2: 5’-AGAATTAGCTCGAATAGAA-3’) or scrambled siRNA control (from Dharmacon) were also used.

### Immunofluorescence

Cells grown on coverslips were fixed with 4% paraformaldehyde and permeabilized using 0.1% saponin containing buffer for 30 mins. Blocking with 1% BSA as well as primary and secondary antibody dilutions were made in permeabilization buffer and incubations were done at 37°C. Images were acquired as Z-stacks using either the confocal or superresolution Airyscan unit of Zeiss 880 confocal microscope using the 63X oil immersion objective with n.a. 1.4 as indicated. Optimal setting as suggested by the software was used to acquire the Z sections. Images are presented as maximum intensity projection or a single Z plane as indicated. Image J software was used for image analysis and Adobe photoshop software was used to assemble the images.

### Electron microscopy

HEK293T cells transfected with GFP or GFP-CPAP or myc-CPAP constructs for 24h were fixed with 2.5% glutaraldehyde in sodium-Cacodylate buffer (Ted Pella Inc) for 30mins and processed as described in^40,70^. After dehydration series with alcohol, cells were embedded in Epoxy resin and cured at 60°C for a couple of days. 70 nm thin sections on copper grids were examined and imaged using the JEOL 1210 transmission electron microscope.

### EGFR internalization assay

HeLa cells were grown in serum free conditions overnight and incubated with media containing cycloheximide [CHX] (5μg/ml) for 1h. Cells were treated in CHX media with EGF for 1h on ice. Cells were washed with chilled serum free media and transferred to 37°C to induce internalization of receptor. For tracking routing of EGF receptor into vesicles by confocal, Alexa Fluor 555 conjugated EGF (250ng/ml) was used. Untagged EGF (10ng/ml) was used for some experiments. Cell surface EGFR expression was determined by staining cells on ice with anti-EGFR-Alexa 647 labeled antibody, followed by acquisition using the FACS verse instrument (BD Biosciences). Data was analyzed using the Flowjo software.

### Recycling assay

Similar to EGF treatment conditions, HeLa cells were grown in serum free conditions overnight and incubated with media containing CHX (5μg/ml) for 1h. Cells were treated in CHX containing media with holotransferrin ligand for 1h on ice. Cells were washed with serum free media and transferred to 37°C to induce internalization of receptor.

### Western blot (WB) and immune blot (IB) assays

Cells were lysed on ice, lysates were spun at 14000 rpm for 20 min, followed by SDS-PAGE, WB and IB. For determining EGFR degradation, cells were collected at indicated time points and lysed using RIPA lysis buffer containing 0.1% SDS and 1% NP-40 detergent. Scanned images of original x-ray films or ChemiDoc images of relevant IB panels are included as supplemental figure 6.

### Image quantification and statistical considerations

Z-stack images were split into single Z planes and red and yellow pixels were quantified to determine the percentage of colocalization. All experiments were performed at least thrice and data from representative experiments has been shown. *P*-values were calculated using GraphPad Prism statistical analysis software.

## Acknowledgements

We would like to thank Dr. T.K. Tang and Dr. Pierre Gonczy for sharing the myc-CPAP/ GFP-CPAP and doxycycline inducible GFP-CPAP expression constructs respectively. We would also like to thank the Cell & Molecular Imaging Shared Resource which is supported by the Hollings Cancer Center, Medical University of South Carolina (P30 CA138313) and the Shared Instrumentation Grant S10 OD018113. We would also like to thank the Flow cytometry and Electron microscopy core facilities at the Medical University of South Carolina. This work was supported by NIH grants R21DE026965 and R21DE026965-02S1 to R.G. and C.V. and MUSC internal funds to C.V.

## Data availability

The datasets generated during and/or analyzed during the current study are available from the corresponding author on reasonable request.

