## Supplemental data for "Centrosomal P4.1-associated protein (CPAP) positively regulates endocytic vesicular transport and lysosome targeting of EGFR"

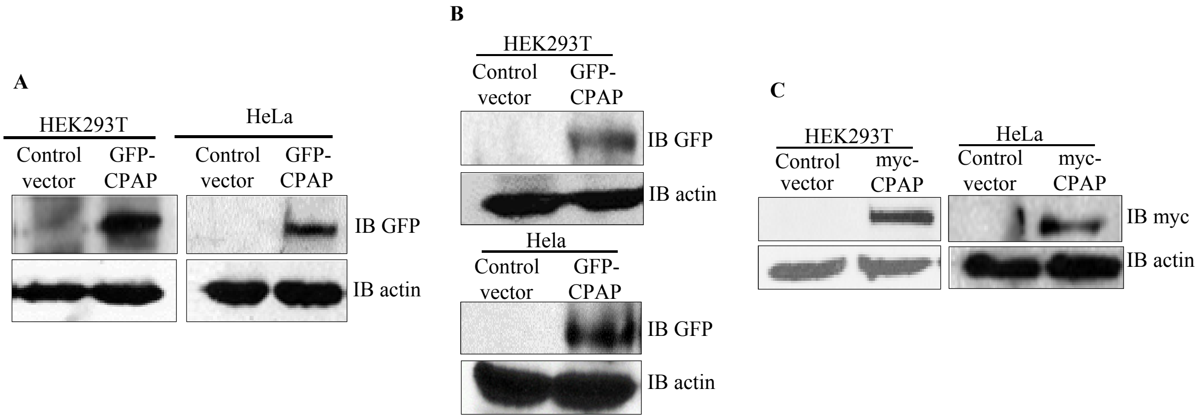


**Supplemental Fig. 1: IBs showing exogenous expression of GFP-CPAP and myc-CPAP.** **A.** HEK293T and HeLa cells were transiently transfected with GFP or GFP-CPAP expression vectors for 24h and subjected to IB to detect GFP-CPAP and β-actin. **B.** HEK293T and HeLa cells expressing GFP or GFP-CPAP under doxycycline (doxy) -inducible promoter were harvested 24h post doxy treatment and subjected to IB to detect GFP-CPAP and β-actin. **C.** HEK293T and HeLa cells were transfected with control or myc-CPAP expression vectors for 24h and subjected to IB to detect myc and β-actin.

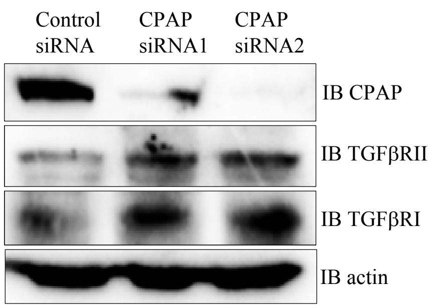


**Supplemental Fig. 2: CPAP depletion causes increased cellular levels of TGF receptors.** HeLa cells were treated with control-siRNA, CPAP-siRNA1 and CPAP-siRNA2 for 72h and subjected to IB to detect CPAP, TGFβRII, TGFβRI and β-actin.

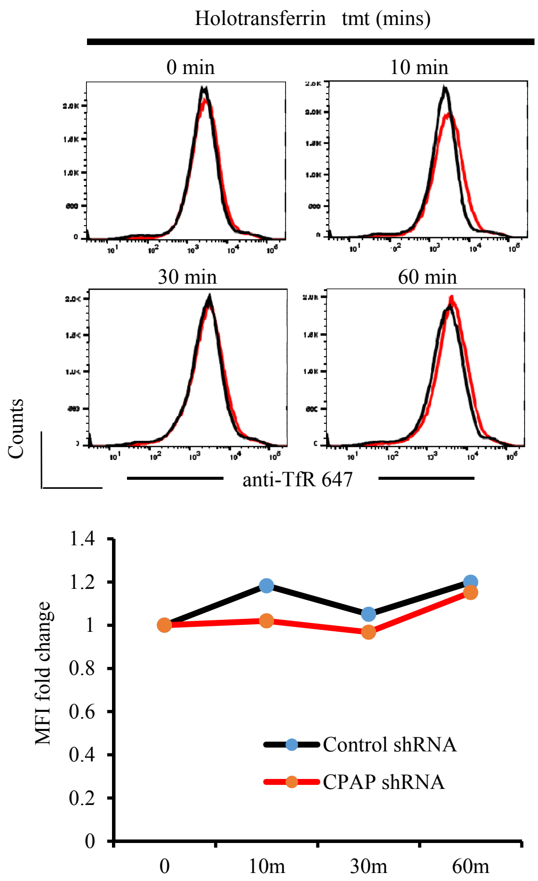


**Supplemental Fig. 3: CPAP depletion does not impact surface levels of TfR.** HeLa cells stably expressing control-shRNA or CPAP-shRNA were subjected to serum starvation overnight, treated with cycloheximide for 1h and holotransferrin, washed and incubated for indicated durations, and subjected to FACS analysis to detect surface levels of TfR after staining using anti-TfR antibody. Representative overlay graphs for each time-point **(upper panel)** and transferrin treatment induced fold changes in TfR specific MFI values, relative to 0 min time-point, (**lower panel**) are shown.

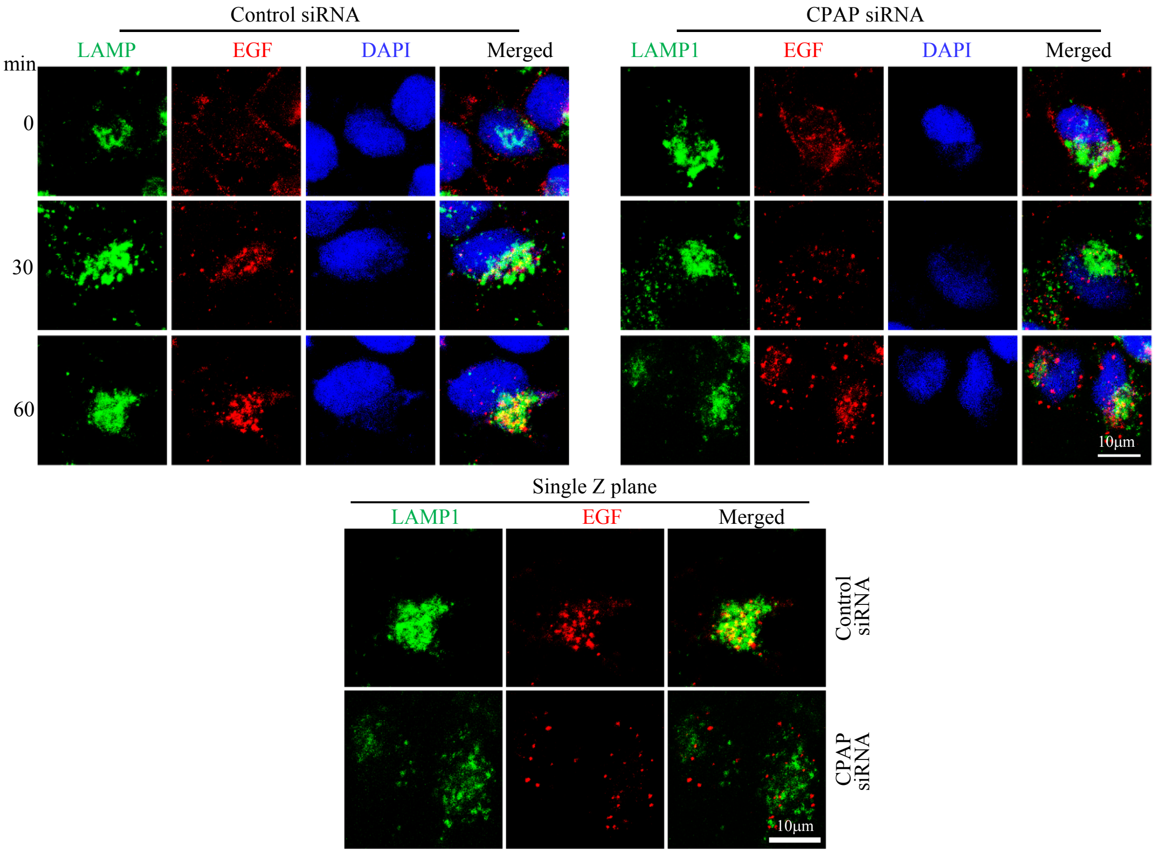


**Supplemental Fig. 4: CPAP depletion results in defective targeting of internalized cell surface receptor to the lysosome.** HeLa cells were treated with control-siRNA or CPAP-siRNA1 for 72h and incubated with Alexa fluor 555-conjugated EGF ligand and left on ice for 1h. Cells were washed with serum free media and transferred to 37°C to initiate receptor internalization. Cells were fixed at indicated time-points, permeabilized and stained for LAMP1 to mark lysosomes. Images were acquired as Z-stacks using Zeiss 880 and representative maximum projection images (upper panel) and single Z plane of relevant images of 60 min time-point (lower panel) are shown.

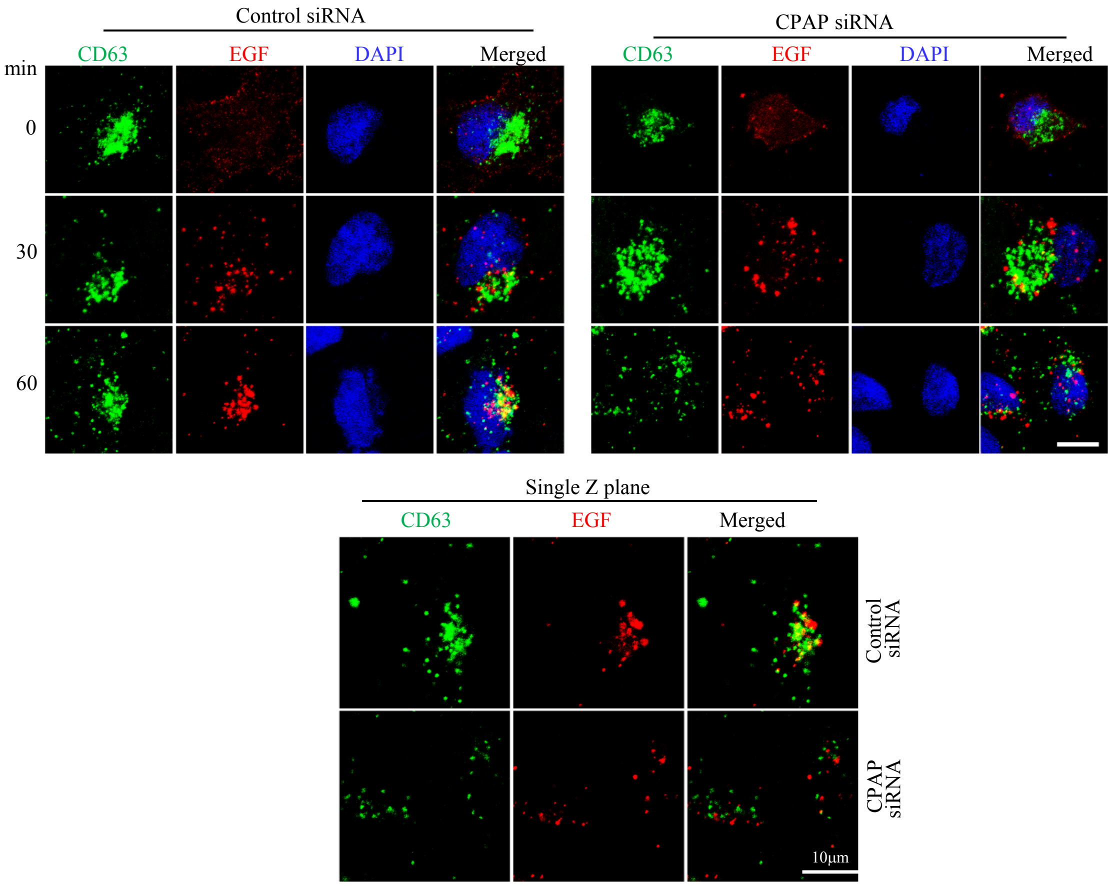


**Supplemental Fig. 5: CPAP depletion results in defective trafficking of internalized cell surface receptor to MVB/late endosome.** HeLa cells were treated with control-siRNA or CPAP-siRNA1 for 72h and incubated with Alexa fluor 555-conjugated EGF ligand and left on ice for 1h. Cells were washed with serum free media and transferred to 37°C to initiate receptor internalization. Cells were fixed at indicated time-points, permeabilized and stained for CD63 to mark MVB/late endosome. Images were acquired as Z-stacks using Zeiss 880 and representative maximum projection images (upper panel) and single Z plane of relevant images of 60 min time-point (lower left panel) are shown.
